# ntEdit: scalable genome sequence polishing

**DOI:** 10.1101/565374

**Authors:** René L Warren, Lauren Coombe, Hamid Mohamadi, Jessica Zhang, Barry Jaquish, Nathalie Isabel, Steven JM Jones, Jean Bousquet, Joerg Bohlmann, Inanç Birol

## Abstract

In the modern genomics era, genome sequence assemblies are routine practice. However, depending on the methodology, resulting drafts may contain considerable base errors. Although utilities exist for genome base polishing, they work best with high read coverage and do not scale well. We developed ntEdit, a Bloom filter-based genome sequence editing utility that scales to large mammalian and conifer genomes.

We first tested ntEdit and the state-of-the-art assembly improvement tools GATK, Pilon and Racon on controlled *E. coli* and *C. elegans* sequence data. Generally, ntEdit performs well at low sequence depths (<20X), fixing the majority (>97%) of base substitutions and indels, and its performance is largely constant with increased coverage. In all experiments conducted using a single CPU, the ntEdit pipeline executed in <14s and <3m, on average, on *E. coli* and *C. elegans*, respectively. We performed similar benchmarks on a sub-20X coverage human genome sequence dataset, inspecting accuracy and resource usage in editing chromosomes 1 and 21, and whole genome. ntEdit scaled linearly, executing in 30-40m on those sequences. We show how ntEdit ran in <2h20m to improve upon long and linked read human genome assemblies of NA12878, using high coverage (54X) Illumina sequence data from the same individual, fixing frame shifts in coding sequences. We also generated 17-fold coverage spruce sequence data from haploid sequence sources (seed megagametophyte), and used it to edit our pseudo haploid assemblies of the 20 Gbp interior and white spruce genomes in <4 and <5h, respectively, making roughly 50M edits at a (substitution+indel) rate of 0.0024.

**Availability:** https://github.com/bcgsc/ntedit

**Supplemental material:** available online.

## Introduction

This past decade, next generation sequencing technologies have greatly improved their throughput. For example, 50-fold coverage of a 20 Gbp conifer genome can today be achieved on a single, 8-lane flowcell of the Illumina HiSeq-X instrument. However, this massive data is creating bottlenecks in bioinformatics pipelines. Typically, with short read data, a pseudo haploid draft genome representing unresolved allelic mixtures results from diploid sequence assembly. Depending on the methodology used, these assemblies may contain appreciable errors. Although many tools exist for base error correction of reads from high throughput sequencing platforms [1], few genome assembly polishing tools are available.

The leading utilities for assembly polishing include GATK [2], Pilon [3] and Racon [4]. Pilon and GATK are well established and comprehensive tools for genome improvements, and include the ability to fill short gaps, fix local misassemblies, and identify and report variant bases. In comparison, Racon is a more recent utility originally designed as a fast nanopore read correction tool. The latter is gaining traction in polishing, using Illumina data, single molecule sequencing (SMS) genome drafts such as those assembled from Pacific Biosciences (PacBio) and Oxford Nanopore (Nanopore) sequence reads [5]. Considered the state-of-the art in polishing accuracy, Pilon is a robust genome assembly improvement tool routinely used to polish microbial and small (<100 Mbp) eukaryotic genomes. It has also been applied to human assemblies [6], but unfortunately scales quadratically in time.

The aforementioned tools all employ read alignments. This paradigm gives context to the bases under scrutiny, albeit at the expense of run time. To address these scalability limitations, we developed ntEdit, a utility that uses words of length k (kmer) for correcting homozygous errors in very large genome (>3Gbp) assemblies. ntEdit employs a succinct Bloom filter data structure for evaluation and correction. By comparing it to the base polishing capabilities of other tools, namely the ability to fix base substitutions and indels, we show how ntEdit produces comparable results, and scales linearly to the large 3-20 Gbp genomes of human and spruce [7].

## Methods

We first run ntHits (v0.0.1; https://github.com/bcgsc/nthits; Supplemental Methods) to remove error kmers from high throughput sequencing data, and build a canonical representation of coverage-thresholded kmers [8] using a Bloom filter, while maintaining a low false positive rate (≈0.0005). The Bloom filter is then read by ntEdit (v1.1.0 with matching kmer length k), and contigs from a supplied assembly are processed in turn (Fig. S1). From each sequence, 5’ to 3’ assembly kmers are used to interrogate the Bloom filter for presence/absence. When a kmer is absent, ntEdit moves the frame over the next (k-1) 3’-end bases along the assembled sequence, and uses a subset of these k kmers to further interrogate the Bloom filter. In its tests, it uses two leniency factors. The first one ensures that only positions that do not have a sufficient subset of k kmer support are edited. The kmers that pass this filter (ie. absent kmers) will have their 3’-end base permuted, in turn, to one of the three alternate bases. Changes are tracked when kmers with the base substitution are contained in the Bloom filter using a kmer threshold set by a second leniency factor. In that case, the remaining alternate base substitution(s) are also investigated. The kmers that do not pass this second filter are inspected for micro-insertion(s) and deletion(s) of up to 5 bases, and changes are tracked as per above. The process repeats until a change is made that has sufficient support, or until all possible edits have been exhausted at that position (Fig. S1). ntEdit outputs a new fasta sequence with the changes applied, and a text file tracking changes made along with with positions. ntEdit is implemented in C++. Detailed methods available online.

## Results and Discussion

We measured the performance of these tools using QUAST [9], comparing simulated genome copies with 0.001 and 0.0001 substitution and indel rates, along with GATK, Pilon, Racon and ntEdit-polished versions to their respective reference genomes (Fig. S2. and Fig. 1). The performance of ntEdit in fixing substitutions and indels was largely constant with increased coverage from 15-50X. It was also similar to the performances of other tools, and consistent even at low (15-20x) read coverages (Fig. 1 panels A, B, E, F, I; Tables S1-S3). Notably, ntEdit had a slightly higher false discovery rate (FDR) under all tested conditions, and generally less than 2 and 1 percent at higher k values (k45 and up), for the larger (3 Gbp) and smaller (<100 Mbp) genomes, respectively. We also note that its FDR diminished with increasing k (Table S4-S6). We determined that, although the Bloom filter false positive rate (FPR) has the potential to affect the corrections if set too high (at FPR >0.001), it did not impact ntEdit’s ability to identify and correct erroneous bases at the FPR levels used in our experiments (Fig. S3, FPR≈0.0005). For *E. coli, C. elegans* and *H. sapiens* the optimum k to use with ntEdit was 25, 35 and 50 (Table S7), respectively. We also tested how iterative ntEdit runs with varied kmer lengths may benefit in correctly fixing additional sites, despite run time trade-offs (Tables S1-S6). Pilon and Racon performed best at higher coverage (≥20X) sequencing data, though they also had higher resource requirements (Fig. 1 panels C, D, G, H, J, K; Tables S1-S3). The ntEdit pipeline ran in 10s, 3m and 38m on average for 15-50X *E. coli* (k25), *C. elegans* (k35) and sub-20X *H. sapiens* (k50) data (Tables S1-S3), respectively. In contrast, Pilon took 4-11m, 2-5h and several hours/days on the same data, and it ran twice as slow at lower (eg. 15X) coverages.

**Fig. 1.**
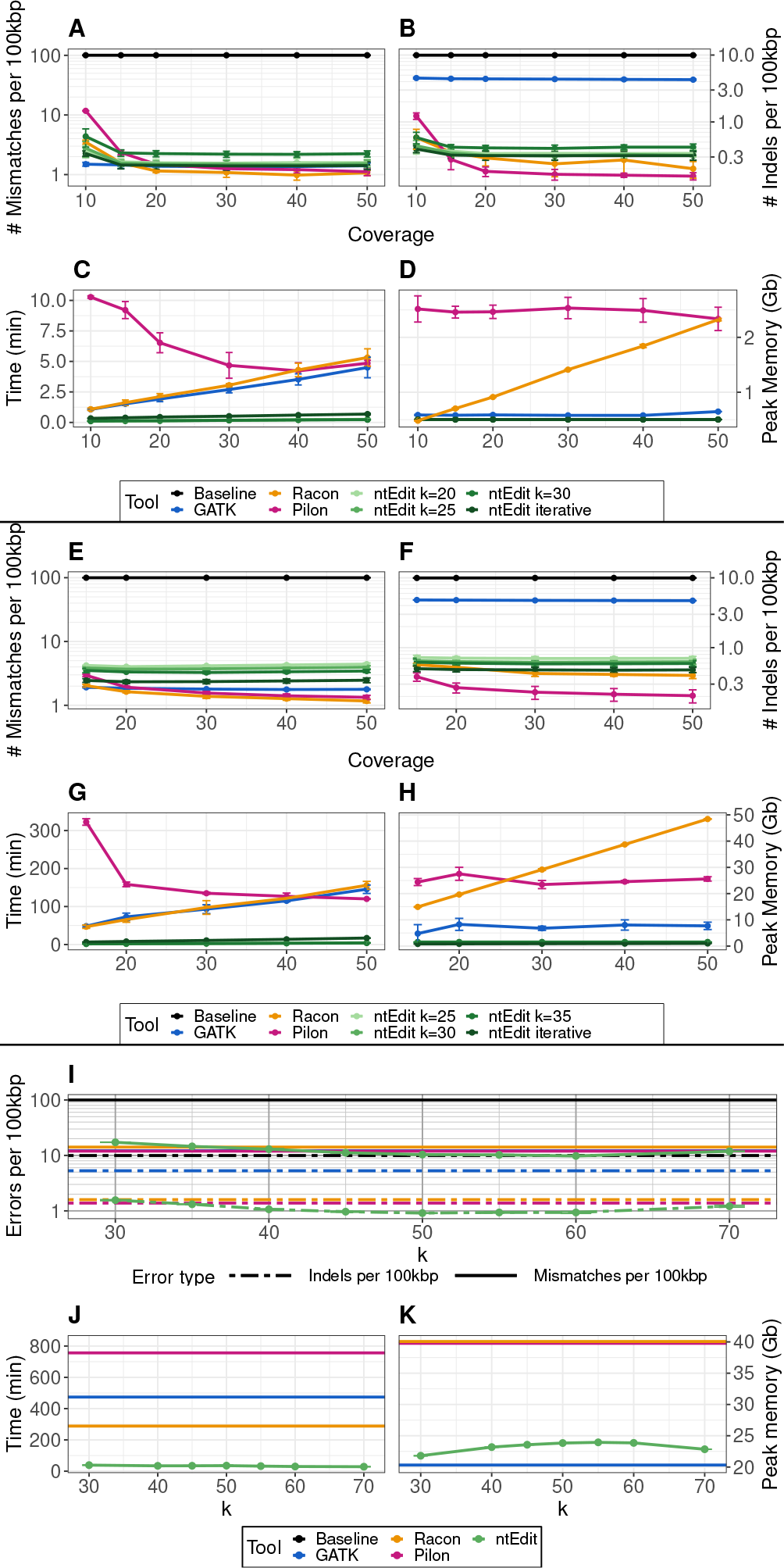
Performance and benchmark of GATK, ntEdit, Pilon and Racon using controlled *E. coli* (panels A, B, C, D) and *C. elegans* (panels E, F, G, H) genomes and *H. sapiens* chromosome 21 (I, J, K) sequences.

Since Pilon executed in >1 week on chromosome 1 (with 48 threads, Table S3), we did not run it on experimental large genome assemblies. Running ntEdit for correction of experimental human NA12878 SMS assemblies with Illumina reads, of which some were already polished [6], led to the recovery of additional complete BUSCO genes [10]. BUSCO measures assembly completeness by mapping single-copy gene products to assemblies. To this effect, a higher number of genes are expected to be recovered when fixing substitutions, but more importantly, when fixing indels causing premature stops and frame shifts. Of the best runs, ntEdit recovered 26 and 366 additional complete BUSCO genes, in Nanopore [11] and PacBio [12] assemblies, respectively (Table S8). As expected given the low indel rate of Illumina data, sequence polishers had only marginal effects on the Supernova assembly using the BUSCO metric, recovering <3 complete BUSCOs. The ntEdit runs on the SMS assemblies were fast, with the whole pipeline (ntHits and ntEdit) executing in 2h, with the ntEdit portion taking two to four minutes at most. The accuracy of GATK and Racon is noteworthy, albeit the former appears to struggle when fixing indels. In order to run Racon in a reasonable amount of time on large (>3Gbp) genomes, we sectioned the drafts (20 sub files), each running in an embarrassingly parallel fashion, with 48 CPU threads. Its cumulative run time was >1.6 days. Without any partitioning of the human genome drafts, Racon runs took over 3 days. On small genomes and less contiguous (N50 length <1Mbp) assemblies, the memory (RAM) usage of ntEdit is largely constant, occupied mainly by the Bloom filter. However, We observe larger memory footprints with more contiguous draft assemblies (Tables S3 and S8).

We also ran ntEdit on the interior (PG29-v4, 20.17 Gbp) and white (*P. glauca* WS77111-v1, 21.90 Gbp) spruce pseudo haploid genome assemblies [13]. Runs took 4h46m and 3h55m in total, with the ntEdit portion running in less than 30 minutes, and required at most 208 GB RAM. Overall 48.3 and 50.5 M haploid edits were made at a substitution and indel rate of 2.3×10^−3^ and 5.0×10^−5^ for the two genomes, respectively (Table 1).

**Table 1.**
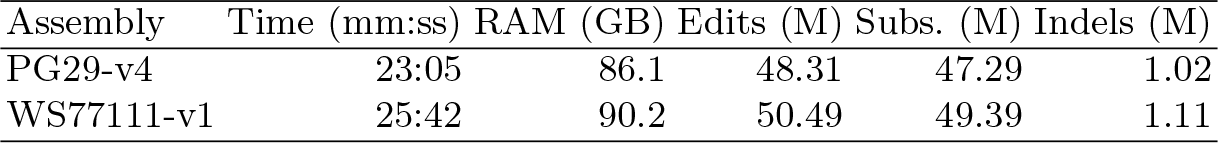
Performance of ntEdit in polishing the pseudo haploid assemblies of interior (PG29-v4) and white (WS77111-v1) spruce, using DNA sequences from their respective haploid megagametophyte tissue. Bloom filter creation with ntHits took 4h23m and 3h29m and required 206.9 GB and 207.8 GB RAM on these data, using 48 CPU threads.

Genome polishing approaches that rely on paired-end read alignments tend to be more accurate overall. This is not surprising since paired-end reads provide more sequence context than kmers. For ntEdit, in its current implementation, additional sequence context is considered up to twice the value of k by exploring a subset of overlapping k kmers having the qualifying change. At the moment, only homozygous errors are targeted, and ntEdit compromises sensitivity in favor of performance. With its fast Bloom filter operations, ntEdit simplifies polishing and “haploidization” of gene and genome sequences since those data structures can be re-used on different sequences, draft assemblies, etc., and in separate polishing runs, without the need to partition the input assembly. We also expect ntEdit to have additional applications in fast mapping of simple nucleotide variations between any two individuals or species’ genomes.

## Acknowledgements

This work was supported by Genome Canada and Genome BC [243FOR, 281ANV]; and the National Institutes of Health [2R01HG007182-04A1]. The content of this
work is solely the responsibility of the authors, and does not necessarily represent the official views of the National Institutes of Health or other funding organizations.

