## Supplemental material for "ntEdit: scalable genome sequence polishing"

### Supplementary Data

#### Supplementary Methods

##### Description of ntHits

ntHits is a streaming algorithm and software tool for *de novo* repeat identification based on the statistical analysis of the kmer content profile of large-scale DNA sequencing data. It first generates the kmer coverage histograms of input datasets using the ntCard algorithm, an efficient streaming algorithm for estimating kmer coverage histograms (Mohamadi et al., 2017). From the obtained kmer coverage histogram, the repetitive kmers would present a long tail to the distribution of kmer coverage profile. ntHits identifies a threshold from the long tail of kmer profile distribution and outputs the highly repetitive kmers with frequency greater than the identified threshold. We have adapted ntHits (v0.0.1) to work with ntEdit by reporting a Bloom filter (--outbloom option) of non-error kmers. On controlled data, we ran ntHits with a set threshold (-c 1 on <20X coverages and –c 2 on ≥ 20X, indicating kmers with a coverage above 1 and 2, respectively). On the experimental human runs, ntHits ran on a 54X coverage Illumina read data in auto mode (nthits options --outbloom --solid -b 36 -k 35 -t 48), using a threshold set by the ntCard algorithm based on the corresponding kmer coverage histogram it computes (Fig. S4).

##### Data and run parameters on controlled experiments

To best assess ntEdit's correction performance we used pIRS v2.0.0 (Hu et al., 2012) to simulate *E. coli*, *C. elegans* and *H. sapiens* triplicate genome copies from their respective *E. coli* K12 MG1655 (U00096.3), *C. elegans* Bristol N2 (NC_003281.10) and *H. sapiens* (GRCh38) reference genomes, with substitution and indel rates of 0.001 and 0.0001, respectively (pirs diploid <genome.fa> -s 0.001 -d 0.0001). These were supplied to ntEdit (-f option) in lieu of draft assemblies (Fig. S2). Using the corresponding reference genomes, we simulated three sets of paired-end 100 bp reads with 0.1% error using Illumina error models at 10-50, 15-50 and 17-fold sequence coverages, respectively (pirs simulate -l 100 -x <coverage> -m 300 -e 0.001 --no-gc-bias --no-indels -B humNew.PE100.matrix.gz). We ran Pilon (v1.22 --changes --diploid --fix snps,indels) and GATK (v4.1.0.0) with bwa mem (v.0.7.17-r1188). GATK also ran with the help of samtools and bcftools (v1.9). The VCF file generated using bcftools was filtered to only include variants with IMF >= 0.7 and (reads supporting variant)/(total reads) >= 0.7 prior to polishing the input assembly with "gatk FastaAlternateReferenceMaker". Racon (v1.3.1), with Minimap2 (v2.15) and ntEdit (v1.1.0 -k 20-70 step 5 or 10, -i 4 -d 5 -x 5 -y 9) with ntHits (v0.0.1 -c 1 --outbloom) also ran on these datasets and the resulting assemblies were assessed with QUAST-LG v5.0.2, comparing the edited copies to their respective reference genome (Fig. S3). To speed up the Racon runs on the full *H. sapiens* genome, following the Minimap2 alignments of the simulated reads to the genome, the assembly, read alignments and reads were partitioned into 24 separate jobs; one per chromosome. Then, Racon polishing of each chromosome was run in parallel on a computing cluster (Intel Zeon E5-2650, 2.20 GHz, 384GB RAM, 48 threads). We also ran ntEdit iteratively on *E. coli*, *C. elegans* and *H. sapiens* at k35, k30, k25 and k40, k35, k30, k25 and k50, k45, k40, respectively.

##### Data and run parameters on experimental data

We polished NA12878 human genome drafts, including an assembly of 1) Pacific Bio sciences (PacBio) (Pendleton et al., 2015), 2) Nanopore (Jain et al., 2018) and 3) 10x Chromium linked (https://support.10xgenomics.com/de-novo-assembly/datasets/2.1.0/wfu) reads with ntEdit (several parameters – see Table S8), Racon (-u -w 500) and GATK (variant filtering same as for controlled experiments) using Illumina reads for correction (downloaded from https://basespace.illumina.com public dataset : HiSeq 2500: TruSeq PCR-Free DNA 2x251 (NA12878), flowcell H00DDBCXX). In order to run Racon in a reasonable amount of time, we first aligned the full read set to each draft assembly with Minimap2, then split the draft assemblies, read alignments and reads into 20 partitions. Racon was run on each partition in parallel on a compute farm (Intel Zeon E5-2650, 2.20 GHz, 384GB RAM), with each job using 48 threads. We assessed the resulting, corrected assemblies, with BUSCO (Simão et al., 2015; v.2.0.1 lineage: euarchontoglires_odb9, eukaryota) to show both completeness and resolution of frameshift errors by reporting recovered gene products. Finally, to show scale, we ran ntEdit (-k50 -i1 -d1 -x5 -y9 -t 48) on the 20-Gbp interior (PG29-v4: Genbank GCA_000411955.5) and white (WS77111-v1: Genbank GCA_000966675.1) spruce genome pseudo haploid assemblies using DNA sequences from their corresponding, haploid tissue source (seed megagametophyte, SRA SRX5086197 and SRX5086196). GATK, ntHits, ntEdit, and Pilon, ran on a single CPU on *E. coli* and *C. elegans*. On controlled and experimental *H. sapiens* data, and spruce, all tools used 48 threads. Run time and memory usage was benchmarked on a CentOS 7 system with 128 CPU Intel(R) Xeon(R) CPU E7-8867 v3 @ 2.50GHz. Additional details are available at : http://www.bcgsc.ca/downloads/btl/ntedit/paper/

#### Supplementary Figures

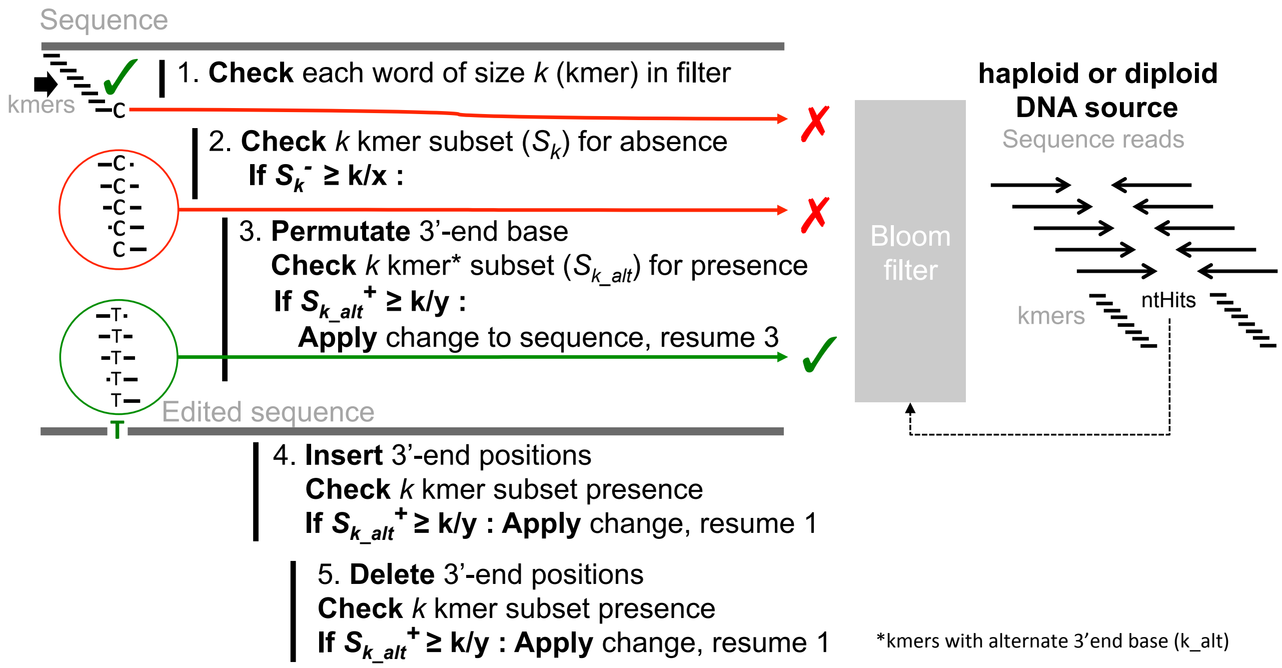


**Supplemental Figure S1. ntEdit approach.** Sequence reads are first shredded into kmers using ntHits, keeping track of kmer multiplicity. The kmers that pass coverage thresholds (ntHits, -c option builds a filter with kmers having a coverage higher than c) are used to construct a Bloom filter (BF). The draft assembly is supplied to ntEdit (-f option, fasta file), along with the BF (-r option) and sequences are read sequentially. Sequence strings are shredded into words of length k (kmers) at a specified value (-k option) matching that used to build the BF, and each kmer from 5' to 3' queries the BF data structure for presence/absence (step 1). When a kmer is not found in the filter, a subset ($S_k$) of overlapping k kmers (defined by k over three, k/3) containing the 3'-end base is queried for absence (step 2). The subset $S_k$, representing a subsampling of k kmers obtained by sliding over 3 bases at a time over k bases, is chosen to minimize the number of checks against the Bloom filter. Of this subset, when the number of **absent (-)** kmers matches or exceeds a threshold defined by $S_k^-$ >= k/x (-x option), representing the majority of kmers in $S_k$, editing takes place (step 3 and beyond), otherwise step 1 resumes. In the former case, the 3'-end base is permuted to one of the three alternate bases (step 3), and the subset ($S_{k\_alt}$) containing the change is assessed for Bloom filter **presence (+)**. When that number matches or exceeds the threshold defined by $S_{k\_alt}^+$ >= k/y (-y option), which means the base substitution qualifies, it is tracked along with the number of supported kmers and the remaining alternate 3'-end base substitutions are also assessed (ie. resuming step 3 until all bases inspected). If the edit does not qualify, then a cycle of base insertion(s) and deletion(s) of up to –i and –d bases begins (step 4, -i option and step 5, -d option, respectively). As is the case for the substitutions, a subset of k kmers containing the indel change is tested for presence. If there are no qualifying changes, then the next alternate 3'-end base is inspected as per above; otherwise the change is applied to the sequence string and the next assembly kmer is inspected (step 1). The process is repeated until a qualifying change or until no suitable edits are found. In the latter case, we go back to step 1. When a change is made, the position on the new sequence is tracked, along with an alternate base with lesser or equal k kmer subset support, when applicable. Currently, ntEdit only tracks cases when edits are made (steps 3-5), and does not flag unedited, missing draft kmers (steps 1-2).

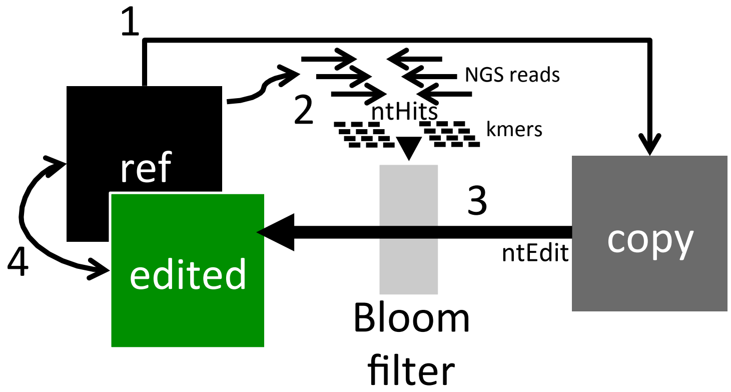


**Supplemental Figure S2. Testing procedure on controlled data.** To test against a ground truth, we made three copies of the *E. coli* K12 MG1655 (U00096.3), *C. elegans* Bristol N2 (NC_003281.10) and *H. sapiens* (GRCh38) reference genomes with pIRS (v2.0.0 pirs diploid <genome.fa> -s 0.001 -d 0.0001), tracking the coordinates and edits made (step 1). Using the reference genomes, we simulated PE100 reads using an Illumina error model at 0.1% error rate, and several fold coverages (10, 15, 20, 30, 40, 50) (pirs simulate -l 100 -x <COVERAGE> -m 300 -e 0.001 --no-gc-bias --no-indels -B humNew.PE100.matrix.gz) and built Bloom filters for each with ntHits (-c 1) (step2). Using the copy, we ran ntEdit/comparators in conjunction with the reference-derived reads (step 3). The resulting draft assembly was compared to the original reference genome and copy with QUAST (v5.02), tracking the number of mismatches and indels per 100kbp (step4) (tables S1-S3). Additionally, we made statistical measures of the performance. We report those values in tables S4, S5, and S6 (below).

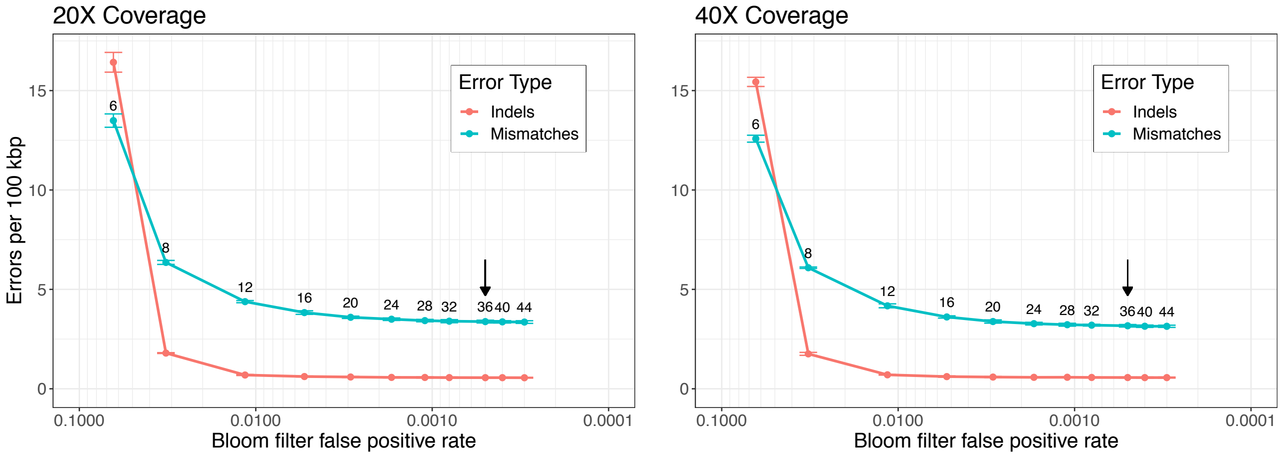


**Supplemental Figure S3. Effect of Bloom filter false positive rate on ntEdits predictions.** On the *C. elegans* controlled experiments (20X and 40X coverage data, left and right panels) and in triplicate experiments, we used ntHits to output Bloom filters (-c 1 --outbloom –k 35), controlling their bit sizes (-b option, values indicated on the plot) but using the same set of reads. The Bloom filter false positive rate (FPR) was calculated as FPR=(pop. count/bit size)$^{hashes}$. We ran ntEdit (-k 35 –i 4 –d 5 –x 5 –y 9) using each Bloom filter as input and the resulting, edited assembly was assessed by QUAST. By default, ntHits outputs Bloom filters with a FPR <0.001 (the ones used in our study were consistently set at b=36, FPR≈0.0005, black arrows). In that range, the Bloom filter FPR has little effect on the performance of ntEdit.

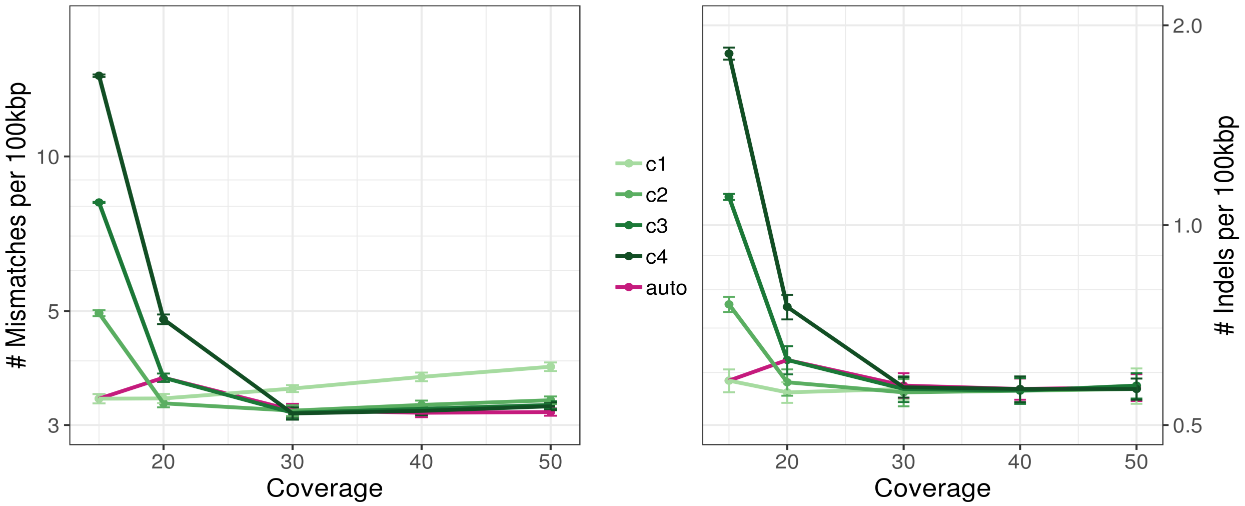


**Supplemental Figure S4. Effect of the ntHits threshold c parameter.** In separate experiment, we ran ntHits using hard thresholds of -c (1,2,3,4) to discard kmers (at those coverage and lower) and build a Bloom filter with the remaining kmers (nthits options -c 1 -b 36 --outbloom -k 35 -t 48). We also ran ntHits in auto mode, where the threshold for discarding erroneous kmers is set automatically by ntCard based on the kmer coverage histogram (nthits options --outbloom --solid -b 36 -k 35 -t 48). We then ran ntEdit using each Bloom filter as input and the resulting edited assembly was assessed with QUAST (ntEdit options -k 35 -d 5 -i 4 –x 5 –y 9). We here show the effect (mismatches and indels per 100kbp, left and right panels, respectively) of setting these thresholds on the *C. elegans* controlled data.

#### Supplemental Tables

**Supplemental table 1. QUAST and resource benchmark results for controlled *E. coli* dataset.**

| Assembly | Cover-age | Mismatches per 100kbp | Indels per 100kbp | Time (min.) | Peak memory (Mb) |
|---|---|---|---|---|---|
| Baseline | N/A | 99.81 +/- 0.85 | 9.99 +/- 0.07 | N/A | N/A |
| GATK | 10 | 1.49 +/- 0.11 | 4.53 +/- 0.13 | 1.07 +/- 0.05 | 599.77 +/- 9.48 |
| | 15 | 1.45 +/- 0.19 | 4.43 +/- 0.12 | 1.52 +/- 0.12 | 595.40 +/- 4.51 |
| | 20 | 1.38 +/- 0.16 | 4.40 +/- 0.06 | 1.92 +/- 0.22 | 601.49 +/- 15.82 |
| | 30 | 1.36 +/- 0.17 | 4.37 +/- 0.05 | 2.70 +/- 0.27 | 593.11 +/- 0.98 |
| | 40 | 1.36 +/- 0.14 | 4.32 +/- 0.13 | 3.53 +/- 0.45 | 592.59 +/- 4.36 |
| | 50 | 1.41 +/- 0.16 | 4.29 +/- 0.10 | 4.50 +/- 0.84 | 663.18 +/- 0.72 |
| Racon | 10 | 3.51 +/- 0.28 | 0.58 +/- 0.19 | 1.08 +/- 0.09 | 494.14 +/- 8.86 |
| | 15 | 1.57 +/- 0.17 | 0.35 +/- 0.05 | 1.62 +/- 0.23 | 720.66 +/- 11.55 |
| | 20 | 1.15 +/- 0.07 | 0.29 +/- 0.07 | 2.11 +/- 0.25 | 934.52 +/- 7.26 |
| | 30 | 1.08 +/- 0.19 | 0.24 +/- 0.09 | 3.05 +/- 0.14 | 1445.02 +/- 7.68 |
| | 40 | 0.98 +/- 0.17 | 0.27 +/- 0.10 | 4.30 +/- 0.55 | 1886.18 +/- 30.53 |
| | 50 | 1.07 +/- 0.11 | 0.20 +/- 0.06 | 5.32 +/- 0.71 | 2378.52 +/- 22.27 |
| Pilon | 10 | 11.73 +/- 0.12 | 1.24 +/- 0.16 | 10.27 +/- 0.10 | 2577.22 +/- 244.04 |
| | 15 | 2.33 +/- 0.25 | 0.24 +/- 0.06 | 9.21 +/- 0.70 | 2518.02 +/- 110.24 |
| | 20 | 1.51 +/- 0.18 | 0.17 +/- 0.02 | 6.53 +/- 0.82 | 2524.06 +/- 124.19 |
| | 30 | 1.26 +/- 0.21 | 0.16 +/- 0.03 | 4.68 +/- 1.06 | 2594.91 +/- 199.33 |
| | 40 | 1.19 +/- 0.18 | 0.16 +/- 0.01 | 4.22 +/- 0.67 | 2550.86 +/- 219.62 |
| | 50 | 1.08 +/- 0.17 | 0.15 +/- 0.02 | 4.85 +/- 0.24 | 2392.18 +/- 217.45 |
| ntEdit k20 | 10 | 2.50 +/- 0.57 | 0.40 +/- 0.07 | 0.11 +/- 0.00 | 515.48 +/- 0.06 |
| | 15 | 1.64 +/- 0.21 | 0.37 +/- 0.05 | 0.13 +/- 0.00 | 515.46 +/- 0.03 |
| | 20 | 1.58 +/- 0.23 | 0.34 +/- 0.03 | 0.14 +/- 0.01 | 515.46 +/- 0.02 |
| | 30 | 1.56 +/- 0.19 | 0.34 +/- 0.04 | 0.19 +/- 0.01 | 515.50 +/- 0.02 |
| | 40 | 1.59 +/- 0.27 | 0.34 +/- 0.04 | 0.22 +/- 0.01 | 515.53 +/- 0.03 |
| | 50 | 1.59 +/- 0.22 | 0.34 +/- 0.04 | 0.26 +/- 0.02 | 515.49 +/- 0.00 |
| ntEdit k25 | 10 | 2.77 +/- 0.83 | 0.44 +/- 0.09 | 0.11 +/- 0.01 | 515.47 +/- 0.03 |
| | 15 | 1.56 +/- 0.19 | 0.34 +/- 0.05 | 0.13 +/- 0.00 | 516.77 +/- 2.30 |
| | 20 | 1.50 +/- 0.18 | 0.32 +/- 0.05 | 0.14 +/- 0.00 | 515.48 +/- 0.00 |
| | 30 | 1.50 +/- 0.25 | 0.31 +/- 0.04 | 0.17 +/- 0.01 | 515.52 +/- 0.04 |
| | 40 | 1.46 +/- 0.18 | 0.33 +/- 0.05 | 0.21 +/- 0.00 | 515.43 +/- 0.06 |
| | 50 | 1.53 +/- 0.22 | 0.32 +/- 0.05 | 0.24 +/- 0.02 | 515.50 +/- 0.01 |
| ntEdit k30 | 10 | 4.36 +/- 1.48 | 0.58 +/- 0.12 | 0.11 +/- 0.01 | 515.48 +/- 0.00 |
| | 15 | 2.31 +/- 0.29 | 0.42 +/- 0.04 | 0.13 +/- 0.01 | 516.80 +/- 2.31 |
| | 20 | 2.25 +/- 0.27 | 0.41 +/- 0.04 | 0.14 +/- 0.01 | 515.47 +/- 0.01 |
| | 30 | 2.20 +/- 0.26 | 0.40 +/- 0.05 | 0.17 +/- 0.01 | 516.85 +/- 2.32 |
| | 40 | 2.19 +/- 0.24 | 0.42 +/- 0.03 | 0.21 +/- 0.01 | 515.43 +/- 0.10 |
| | 50 | 2.23 +/- 0.27 | 0.42 +/- 0.05 | 0.23 +/- 0.02 | 515.47 +/- 0.04 |
| ntEdit k35,k30, k25 | 10 | 2.26 +/- 0.23 | 0.39 +/- 0.05 | 0.34 +/- 0.01 | 516.84 +/- 2.30 |
| | 15 | 1.45 +/- 0.20 | 0.32 +/- 0.03 | 0.39 +/- 0.02 | 515.48 +/- 0.03 |
| | 20 | 1.43 +/- 0.21 | 0.31 +/- 0.05 | 0.44 +/- 0.06 | 515.50 +/- 0.03 |
| | 30 | 1.40 +/- 0.21 | 0.31 +/- 0.05 | 0.51 +/- 0.00 | 515.51 +/- 0.03 |
| | 40 | 1.40 +/- 0.21 | 0.31 +/- 0.05 | 0.60 +/- 0.01 | 515.51 +/- 0.05 |
| | 50 | 1.42 +/- 0.22 | 0.31 +/- 0.05 | 0.69 +/- 0.06 | 515.45 +/- 0.07 |

**Supplemental table 2. QUAST and resource benchmark results for controlled *C. elegans* dataset.**

| Assembly | Cover-age | Mismatches per 100 kbp | Indels per 100 kbp | Time (min.) | Peak memory (Mb) |
|---|---|---|---|---|---|
| Baseline | N/A | 99.82 +/- 0.15 | 9.91 +/- 0.05 | N/A | N/A |
| GATK | 15 | 1.91 +/- 0.04 | 4.81 +/- 0.01 | 48.40 +/- 2.47 | 4878.71 +/- 3481.10 |
| | 20 | 1.85 +/- 0.04 | 4.79 +/- 0.01 | 72.84 +/- 9.83 | 8473.77 +/- 2345.65 |
| | 30 | 1.80 +/- 0.05 | 4.75 +/- 0.03 | 93.13 +/- 11.65 | 6949.83 +/- 816.23 |
| | 40 | 1.77 +/- 0.05 | 4.73 +/- 0.01 | 114.98 +/- 2.50 | 8236.92 +/- 2000.48 |
| | 50 | 1.78 +/- 0.04 | 4.71 +/- 0.03 | 145.52 +/- 11.03 | 7894.65 +/- 1434.75 |
| Pilon | 15 | 2.90 +/- 0.07 | 0.36 +/- 0.01 | 322.63 +/- 8.63 | 24986.26 +/- 1367.40 |
| | 20 | 1.89 +/- 0.05 | 0.25 +/- 0.02 | 158.40 +/- 6.10 | 28191.86 +/- 2538.10 |
| | 30 | 1.52 +/- 0.08 | 0.21 +/- 0.02 | 134.97 +/- 2.62 | 24005.51 +/- 1584.47 |
| | 40 | 1.38 +/- 0.09 | 0.19 +/- 0.01 | 126.87 +/- 8.55 | 25133.68 +/- 468.06 |
| | 50 | 1.30 +/- 0.09 | 0.18 +/- 0.01 | 120.07 +/- 0.85 | 26211.87 +/- 853.61 |
| Racon | 15 | 2.03 +/- 0.10 | 0.57 +/- 0.03 | 46.87 +/- 4.40 | 15288.99 +/- 133.17 |
| | 20 | 1.63 +/- 0.07 | 0.52 +/- 0.04 | 65.53 +/- 6.97 | 20127.11 +/- 109.80 |
| | 30 | 1.38 +/- 0.10 | 0.43 +/- 0.04 | 97.48 +/- 17.99 | 29838.36 +/- 336.28 |
| | 40 | 1.27 +/- 0.08 | 0.41 +/- 0.03 | 121.61 +/- 5.47 | 39658.80 +/- 139.50 |
| | 50 | 1.17 +/- 0.08 | 0.40 +/- 0.04 | 155.86 +/- 10.39 | 49578.65 +/- 131.40 |
| ntEdit k25 | 15 | 4.22 +/- 0.04 | 0.72 +/- 0.05 | 1.87 +/- 0.15 | 1563.39 +/- 1.18 |
| | 20 | 4.04 +/- 0.12 | 0.71 +/- 0.04 | 2.14 +/- 0.06 | 1561.03 +/- 0.58 |
| | 30 | 4.17 +/- 0.13 | 0.70 +/- 0.05 | 2.92 +/- 0.16 | 1562.24 +/- 1.11 |
| | 40 | 4.29 +/- 0.17 | 0.70 +/- 0.04 | 3.58 +/- 0.44 | 1562.66 +/- 1.54 |
| | 50 | 4.42 +/- 0.19 | 0.70 +/- 0.05 | 4.54 +/- 0.35 | 1563.29 +/- 1.34 |
| ntEdit k30 | 15 | 3.85 +/- 0.15 | 0.65 +/- 0.07 | 1.63 +/- 0.09 | 1567.38 +/- 1.03 |
| | 20 | 3.69 +/- 0.13 | 0.63 +/- 0.05 | 2.12 +/- 0.08 | 1565.45 +/- 0.49 |
| | 30 | 3.73 +/- 0.17 | 0.62 +/- 0.06 | 2.71 +/- 0.11 | 1567.07 +/- 0.78 |
| | 40 | 3.84 +/- 0.19 | 0.62 +/- 0.05 | 3.68 +/- 0.07 | 1566.81 +/- 1.59 |
| | 50 | 3.96 +/- 0.21 | 0.63 +/- 0.05 | 4.46 +/- 0.05 | 1568.74 +/- 0.47 |
| ntEdit k35 | 15 | 3.52 +/- 0.28 | 0.62 +/- 0.07 | 1.61 +/- 0.02 | 1570.03 +/- 0.90 |
| | 20 | 3.34 +/- 0.08 | 0.60 +/- 0.05 | 1.99 +/- 0.12 | 1567.67 +/- 0.27 |
| | 30 | 3.27 +/- 0.15 | 0.58 +/- 0.05 | 2.67 +/- 0.19 | 1568.75 +/- 2.27 |
| | 40 | 3.35 +/- 0.15 | 0.59 +/- 0.05 | 3.34 +/- 0.14 | 1569.53 +/- 0.26 |
| | 50 | 3.44 +/- 0.19 | 0.59 +/- 0.05 | 4.10 +/- 0.25 | 1569.12 +/- 1.65 |
| ntEdit k40 | 15 | 4.09 +/- 0.48 | 0.66 +/- 0.08 | 1.54 +/- 0.04 | 1571.28 +/- 1.63 |
| | 20 | 3.91 +/- 0.08 | 0.64 +/- 0.03 | 1.88 +/- 0.08 | 1569.42 +/- 1.53 |
| | 30 | 3.62 +/- 0.16 | 0.60 +/- 0.04 | 2.50 +/- 0.14 | 1570.14 +/- 0.44 |
| | 40 | 3.69 +/- 0.15 | 0.61 +/- 0.05 | 3.21 +/- 0.16 | 1570.79 +/- 1.84 |
| | 50 | 3.77 +/- 0.18 | 0.61 +/- 0.05 | 3.99 +/- 0.28 | 1571.92 +/- 0.52 |
| ntEdit k50 | 15 | 5.53 +/- 1.31 | 0.79 +/- 0.15 | 1.46 +/- 0.08 | 1570.54 +/- 2.00 |
| | 20 | 5.48 +/- 0.53 | 0.79 +/- 0.04 | 1.95 +/- 0.02 | 1570.67 +/- 1.57 |
| | 30 | 4.00 +/- 0.13 | 0.65 +/- 0.05 | 2.43 +/- 0.02 | 1573.87 +/- 1.25 |
| | 40 | 3.98 +/- 0.17 | 0.64 +/- 0.05 | 2.89 +/- 0.13 | 1574.05 +/- 1.45 |
| | 50 | 4.01 +/- 0.16 | 0.65 +/- 0.05 | 3.65 +/- 0.28 | 1572.71 +/- 0.55 |
| ntEdit k40,k35, k30,k25 | 15 | 2.45 +/- 0.17 | 0.50 +/- 0.05 | 6.55 +/- 0.26 | 844.53 +/- 0.12 |
| | 20 | 2.34 +/- 0.15 | 0.48 +/- 0.04 | 8.03 +/- 0.16 | 870.45 +/- 0.09 |
| | 30 | 2.35 +/- 0.18 | 0.48 +/- 0.04 | 10.76 +/- 0.19 | 926.35 +/- 0.18 |
| | 40 | 2.41 +/- 0.19 | 0.47 +/- 0.04 | 13.69 +/- 0.42 | 982.85 +/- 0.07 |
| | 50 | 2.47 +/- 0.22 | 0.48 +/- 0.04 | 17.03 +/- 0.76 | 1039.89 +/- 0.49 |

**Supplemental table 3. QUAST and resource benchmark results for 17-fold coverage controlled *H. sapiens* dataset.**

| Assembly | | Mismatches per 100 kbp | Indels per 100 kbp | Time (min.) | Peak Memory (GB) |
|---|---|---|---|---|---|
| *Chromosome 1* | | | | | |
| Baseline | | 99.84 +/- 0.02 | 10.08 +/- 0.00 | N/A | N/A |
| GATK | | 3.60 +/- 0.04 | 4.84 +/- 0.05 | 515.82 +/- 4.66 | 20.32 +/- 0.27 |
| Pilon | | 3.81 +/- 0.02 | 0.48 +/- 0.01 | 11854.90 +/- 2121.66 | 170.90 +/- 20.92 |
| Racon | | 5.12 +/- 0.11 | 0.66 +/- 0.06 | 337.32 +/- 39.33 | 40.06 +/- 0.00 |
| ntEdit | k30 | 16.26 +/- 0.24 | 1.64 +/- 0.03 | 43.07 +/- 3.84 | 24.02 +/- 0.00 |
| | k40 | 11.53 +/- 0.08 | 1.11 +/- 0.02 | 36.28 +/- 5.77 | 24.56 +/- 0.00 |
| | k45 | 9.49 +/- 0.07 | 0.96 +/- 0.02 | 37.21 +/- 5.11 | 24.72 +/- 0.00 |
| | k50 | 8.62 +/- 0.10 | 0.91 +/- 0.02 | 38.39 +/- 8.82 | 24.67 +/- 0.30 |
| | k55 | 8.50 +/- 0.09 | 0.91 +/- 0.02 | 31.91 +/- 2.40 | 24.90 +/- 0.00 |
| | k60 | 8.31 +/- 0.08 | 0.93 +/- 0.02 | 29.55 +/- 1.31 | 24.89 +/- 0.00 |
| | k50,45, 40 | 8.05 +/- 0.08 | 0.85 +/- 0.02 | 102.25 +/- 9.10 | 23.84 +/- 0.00 |
| *Chromosome 21* | | | | | |
| Baseline | | 99.96 +/- 0.30 | 9.96 +/- 0.06 | N/A | N/A |
| GATK | | 12.15 +/- 0.17 | 5.29 +/- 0.12 | 473.81 +/- 8.76 | 20.32 +/- 0.27 |
| Pilon | | 11.95 +/- 0.16 | 1.38 +/- 0.01 | 756.28 +/- 72.92 | 39.77 +/- 0.20 |
| Racon | | 14.13 +/- 0.09 | 1.59 +/- 0.04 | 288.33 +/- 36.80 | 40.06 +/- 0.00 |
| ntEdit | k30 | 17.24 +/- 0.05 | 1.56 +/- 0.03 | 38.66 +/- 1.59 | 21.81 +/- 0.00 |
| | k40 | 12.97 +/- 0.28 | 1.07 +/- 0.02 | 34.17 +/- 1.42 | 23.19 +/- 0.00 |
| | k45 | 11.18 +/- 0.06 | 0.97 +/- 0.02 | 34.48 +/- 0.47 | 23.57 +/- 0.00 |
| | k50 | 10.37 +/- 0.19 | 0.91 +/- 0.01 | 35.40 +/- 3.43 | 23.84 +/- 0.00 |
| | k55 | 10.22 +/- 0.18 | 0.94 +/- 0.03 | 32.32 +/- 2.63 | 23.95 +/- 0.00 |
| | k60 | 9.78 +/- 0.17 | 0.94 +/- 0.04 | 29.81 +/- 1.33 | 23.86 +/- 0.00 |
| | k70 | 11.87 +/- 0.12 | 1.21 +/- 0.05 | 28.48 +/- 0.03 | 22.84 +/- 0.00 |
| | k50,45, 40 | 9.83 +/- 0.17 | 0.83 +/- 0.01 | 96.90 +/- 4.67 | 23.84 +/- 0.00 |
| *Genome* | | | | | |
| Baseline | | 100.00 +/- 0.03 | 10.01 +/- 0.01 | N/A | N/A |
| GATK | | 3.64 +/- 0.01 | 4.83 +/- 0.01 | 1086.93 +/- 10.12 | 20.32 +/- 0.27 |
| Pilon | | N/A | N/A | N/A | N/A |
| Racon* | | 5.16 +/- 0.06 | 0.60 +/- 0.01 | 1455.60 +/- 131.74 | 40.06 +/- 0.00 |
| ntEdit | k40 | 12.07 +/- 0.04 | 1.08 +/- 0.00 | 39.99 +/- 9.65 | 147.10 +/- 24.62 |
| | k45 | 10.16 +/- 0.02 | 0.95 +/- 0.00 | 46.25 +/- 6.11 | 129.80 +/- 5.26 |
| | k50 | 9.46 +/- 0.03 | 0.91 +/- 0.00 | 38.79 +/- 9.89 | 149.28 +/- 25.15 |
| | k60 | 9.47 +/- 0.02 | 0.96 +/- 0.01 | 32.06 +/- 2.41 | 154.79 +/- 18.86 |

* Racon polishing on the full simulated human genome assembly was performed on each chromosome individually in parallel, and the runtimes were summed.

**Supplemental table 4. Performance of polishing utilities on fixing substitutions and indels on simulated *E. coli* genome copies.**

| Tool | Cov. | k | Sensitivity | Precision | FDR | FPR | Specificity | F1 |
|---|---|---|---|---|---|---|---|---|
| ntEdit | 10 | 20 | 0.9711 | 0.9938 | 0.0062 | 0.0011 | 0.9989 | 0.9823 |
| | | 25 | 0.9691 | 0.9933 | 0.0067 | 0.0011 | 0.9989 | 0.9811 |
| | | 30 | 0.9572 | 0.9944 | 0.0056 | 0.0010 | 0.9990 | 0.9754 |
| | | 35 | 0.9497 | 0.9945 | 0.0055 | 0.0010 | 0.9990 | 0.9716 |
| | | 40 | 0.9317 | 0.9948 | 0.0052 | 0.0010 | 0.9990 | 0.9622 |
| | | 35,30,25 | 0.9708 | 0.9940 | 0.0060 | 0.0011 | 0.9989 | 0.9823 |
| GATK | | N/A | 0.9785 | 0.9944 | 0.0056 | 0.0011 | 0.9989 | 0.9863 |
| Pilon | | N/A | 0.8780 | 0.9940 | 0.0060 | 0.0010 | 0.9990 | 0.9324 |
| ntEdit | 15 | 20 | 0.9762 | 0.9938 | 0.0062 | 0.0011 | 0.9989 | 0.9849 |
| | | 25 | 0.9769 | 0.9934 | 0.0066 | 0.0011 | 0.9989 | 0.9850 |
| | | 30 | 0.9699 | 0.9943 | 0.0057 | 0.0011 | 0.9989 | 0.9820 |
| | | 35 | 0.9689 | 0.9942 | 0.0058 | 0.0011 | 0.9989 | 0.9814 |
| | | 40 | 0.9612 | 0.9945 | 0.0055 | 0.0011 | 0.9989 | 0.9776 |
| | | 35,30,25 | 0.9782 | 0.9940 | 0.0060 | 0.0011 | 0.9989 | 0.9860 |
| GATK | | N/A | 0.9800 | 0.9944 | 0.0056 | 0.0011 | 0.9989 | 0.9872 |
| Pilon | | N/A | 0.9710 | 0.9940 | 0.0060 | 0.0011 | 0.9989 | 0.9824 |
| ntEdit | 20 | 20 | 0.9765 | 0.9939 | 0.0061 | 0.0011 | 0.9989 | 0.9851 |
| | | 25 | 0.9770 | 0.9934 | 0.0066 | 0.0011 | 0.9989 | 0.9851 |
| | | 30 | 0.9702 | 0.9943 | 0.0057 | 0.0011 | 0.9989 | 0.9821 |
| | | 35 | 0.9690 | 0.9943 | 0.0057 | 0.0011 | 0.9989 | 0.9815 |
| | | 40 | 0.9599 | 0.9945 | 0.0055 | 0.0011 | 0.9989 | 0.9769 |
| | | 35,30,25 | 0.9785 | 0.9941 | 0.0059 | 0.0011 | 0.9989 | 0.9862 |
| GATK | | N/A | 0.9813 | 0.9942 | 0.0058 | 0.0011 | 0.9989 | 0.9877 |
| Pilon | | N/A | 0.9790 | 0.9939 | 0.0061 | 0.0011 | 0.9989 | 0.9863 |
| ntEdit | 30 | 20 | 0.9765 | 0.9938 | 0.0062 | 0.0011 | 0.9989 | 0.9851 |
| | | 25 | 0.9769 | 0.9934 | 0.0066 | 0.0011 | 0.9989 | 0.9851 |
| | | 30 | 0.9706 | 0.9943 | 0.0057 | 0.0011 | 0.9989 | 0.9823 |
| | | 35 | 0.9703 | 0.9943 | 0.0057 | 0.0011 | 0.9989 | 0.9822 |
| | | 40 | 0.9643 | 0.9945 | 0.0055 | 0.0011 | 0.9989 | 0.9792 |
| | | 35,30,25 | 0.9787 | 0.9941 | 0.0059 | 0.0011 | 0.9989 | 0.9864 |
| GATK | | N/A | 0.9820 | 0.9940 | 0.0060 | 0.0011 | 0.9989 | 0.9880 |
| Pilon | | N/A | 0.9811 | 0.9939 | 0.0061 | 0.0011 | 0.9989 | 0.9875 |
| ntEdit | 40 | 20 | 0.9763 | 0.9938 | 0.0062 | 0.0011 | 0.9989 | 0.9849 |
| | | 25 | 0.9773 | 0.9934 | 0.0066 | 0.0011 | 0.9989 | 0.9853 |
| | | 30 | 0.9705 | 0.9943 | 0.0057 | 0.0011 | 0.9989 | 0.9823 |
| | | 35 | 0.9705 | 0.9943 | 0.0057 | 0.0011 | 0.9989 | 0.9823 |
| | | 40 | 0.9640 | 0.9945 | 0.0055 | 0.0011 | 0.9989 | 0.9790 |
| | | 35,30,25 | 0.9787 | 0.9941 | 0.0059 | 0.0011 | 0.9989 | 0.9864 |
| GATK | | N/A | 0.9821 | 0.9942 | 0.0058 | 0.0011 | 0.9989 | 0.9881 |
| Pilon | | N/A | 0.9817 | 0.9938 | 0.0062 | 0.0011 | 0.9989 | 0.9877 |
| ntEdit | 50 | 20 | 0.9765 | 0.9939 | 0.0061 | 0.0011 | 0.9989 | 0.9851 |
| | | 25 | 0.9766 | 0.9933 | 0.0067 | 0.0011 | 0.9989 | 0.9849 |
| | | 30 | 0.9700 | 0.9943 | 0.0057 | 0.0011 | 0.9989 | 0.9821 |
| | | 35 | 0.9704 | 0.9943 | 0.0057 | 0.0011 | 0.9989 | 0.9822 |
| | | 40 | 0.9640 | 0.9945 | 0.0055 | 0.0011 | 0.9989 | 0.9790 |
| | | 35,30,25 | 0.9787 | 0.9940 | 0.0060 | 0.0011 | 0.9989 | 0.9863 |
| GATK | | N/A | 0.9828 | 0.9942 | 0.0058 | 0.0011 | 0.9989 | 0.9885 |
| Pilon | | N/A | 0.9827 | 0.9939 | 0.0061 | 0.0011 | 0.9989 | 0.9883 |

*NOTE: Mean over 3 runs is shown for all statistics. Racon stats are unavailable as it doesn’t output base edit coordinates

**Supplemental table 5. Performance of polishing utilities on fixing substitutions and indels on simulated *C. elegans* genome copies.**

| Tool | Coverage | k | Sensitivity | Precision | FDR | FPR | Specificity | F1 |
|---|---|---|---|---|---|---|---|---|
| ntEdit | 15 | 25 | 0.9543 | 0.9866 | 0.0134 | 0.0010 | 0.9990 | 0.9702 |
| | | 30 | 0.9569 | 0.9917 | 0.0083 | 0.0011 | 0.9989 | 0.9740 |
| | | 35 | 0.9603 | 0.9922 | 0.0078 | 0.0011 | 0.9989 | 0.9760 |
| | | 40 | 0.9552 | 0.9929 | 0.0071 | 0.0010 | 0.9990 | 0.9737 |
| | | 50 | 0.9447 | 0.9935 | 0.0065 | 0.0010 | 0.9990 | 0.9685 |
| | | 40,35, 30,25 | 0.9706 | 0.9921 | 0.0079 | 0.0011 | 0.9989 | 0.9812 |
| GATK | | N/A | 0.9746 | 0.9938 | 0.0062 | 0.0011 | 0.9989 | 0.9841 |
| Pilon | | N/A | 0.9648 | 0.9937 | 0.0063 | 0.0011 | 0.9989 | 0.9791 |
| ntEdit | 20 | 25 | 0.9561 | 0.9869 | 0.0131 | 0.0010 | 0.9990 | 0.9713 |
| | | 30 | 0.9580 | 0.9919 | 0.0081 | 0.0011 | 0.9989 | 0.9747 |
| | | 35 | 0.9608 | 0.9923 | 0.0077 | 0.0011 | 0.9989 | 0.9762 |
| | | 40 | 0.9545 | 0.9930 | 0.0070 | 0.0010 | 0.9990 | 0.9734 |
| | | 50 | 0.9361 | 0.9935 | 0.0065 | 0.0010 | 0.9990 | 0.9640 |
| | | 40,35, 30,25 | 0.9715 | 0.9922 | 0.0078 | 0.0011 | 0.9989 | 0.9817 |
| GATK | | N/A | 0.9756 | 0.9937 | 0.0063 | 0.0011 | 0.9989 | 0.9846 |
| Pilon | | N/A | 0.9746 | 0.9937 | 0.0063 | 0.0011 | 0.9989 | 0.9840 |
| ntEdit | 30 | 25 | 0.9552 | 0.9869 | 0.0131 | 0.0010 | 0.9990 | 0.9708 |
| | | 30 | 0.9580 | 0.9919 | 0.0081 | 0.0011 | 0.9989 | 0.9747 |
| | | 35 | 0.9619 | 0.9923 | 0.0077 | 0.0011 | 0.9989 | 0.9769 |
| | | 40 | 0.9581 | 0.9930 | 0.0070 | 0.0011 | 0.9989 | 0.9752 |
| | | 50 | 0.9536 | 0.9936 | 0.0064 | 0.0010 | 0.9990 | 0.9732 |
| | | 40,35, 30,25 | 0.9715 | 0.9922 | 0.0078 | 0.0011 | 0.9989 | 0.9817 |
| GATK | | N/A | 0.9767 | 0.9936 | 0.0064 | 0.0011 | 0.9989 | 0.9850 |
| Pilon | | N/A | 0.9782 | 0.9937 | 0.0063 | 0.0011 | 0.9989 | 0.9859 |
| ntEdit | 40 | 25 | 0.9542 | 0.9869 | 0.0131 | 0.0010 | 0.9990 | 0.9702 |
| | | 30 | 0.9571 | 0.9918 | 0.0082 | 0.0011 | 0.9989 | 0.9742 |
| | | 35 | 0.9612 | 0.9922 | 0.0078 | 0.0011 | 0.9989 | 0.9764 |
| | | 40 | 0.9574 | 0.9930 | 0.0070 | 0.0011 | 0.9989 | 0.9749 |
| | | 50 | 0.9541 | 0.9936 | 0.0064 | 0.0010 | 0.9990 | 0.9735 |
| ntEdit | | 40,35, 30,25 | 0.9711 | 0.9921 | 0.0079 | 0.0011 | 0.9989 | 0.9815 |
| GATK | | N/A | 0.9774 | 0.9936 | 0.0064 | 0.0011 | 0.9989 | 0.9854 |
| Pilon | | N/A | 0.9796 | 0.9937 | 0.0063 | 0.0011 | 0.9989 | 0.9866 |
| ntEdit | 50 | 25 | 0.9531 | 0.9867 | 0.0133 | 0.0010 | 0.9990 | 0.9696 |
| | | 30 | 0.9563 | 0.9918 | 0.0082 | 0.0010 | 0.9990 | 0.9737 |
| | | 35 | 0.9606 | 0.9923 | 0.0077 | 0.0011 | 0.9989 | 0.9761 |
| | | 40 | 0.9569 | 0.9930 | 0.0070 | 0.0011 | 0.9989 | 0.9746 |
| | | 50 | 0.9538 | 0.9936 | 0.0064 | 0.0010 | 0.9990 | 0.9733 |
| | | 40,35, 30,25 | 0.9708 | 0.9921 | 0.0079 | 0.0011 | 0.9989 | 0.9813 |
| GATK | | N/A | 0.9608 | 0.9926 | 0.0074 | 0.0011 | 0.9989 | 0.9763 |
| Pilon | | N/A | 0.9803 | 0.9938 | 0.0062 | 0.0011 | 0.9989 | 0.9870 |

*NOTE: Mean over 3 runs is shown for all statistics. Racon stats are unavailable, as it does not output base edit coordinates

**Supplemental table 6. Performance of polishing utilities on fixing substitution and indel on simulated *H. sapiens* chromosomes 1 and 21 and genome copies using 17-fold coverage sequence reads.**

| Tool | k | Sensitivity | Precision | FDR | FPR | Specificity | F1 |
|---|---|---|---|---|---|---|---|
| *Chromosome 1* | | | | | | | |
| ntEdit | 30 | 0.8486 | 0.9770 | 0.0230 | 0.0009 | 0.9991 | 0.9083 |
| | 35 | 0.8763 | 0.9780 | 0.0220 | 0.0009 | 0.9991 | 0.9244 |
| | 40 | 0.8899 | 0.9817 | 0.0183 | 0.0009 | 0.9991 | 0.9335 |
| | 45 | 0.9055 | 0.9836 | 0.0164 | 0.0009 | 0.9991 | 0.9429 |
| | 50 | 0.9106 | 0.9857 | 0.0143 | 0.0009 | 0.9991 | 0.9466 |
| | 55 | 0.9099 | 0.9869 | 0.0131 | 0.0009 | 0.9991 | 0.9468 |
| | 60 | 0.9100 | 0.9877 | 0.0123 | 0.0009 | 0.9991 | 0.9473 |
| | 70 | 0.8737 | 0.9885 | 0.0115 | 0.0009 | 0.9991 | 0.9276 |
| | 50,45,40 | 0.9184 | 0.9841 | 0.0159 | 0.0009 | 0.9991 | 0.9501 |
| GATK | N/A | 0.9349 | 0.9790 | 0.0210 | 0.0009 | 0.9991 | 0.9564 |
| Pilon | N/A | 0.9491 | 0.9907 | 0.0093 | 0.0010 | 0.9990 | 0.9695 |
| Racon | N/A | N/A | N/A | N/A | N/A | N/A | N/A |
| *Chromosome 21* | | | | | | | |
| ntEdit | 30 | 0.8381 | 0.9770 | 0.0230 | 0.0008 | 0.9992 | 0.9023 |
| | 35 | 0.8643 | 0.9775 | 0.0225 | 0.0008 | 0.9992 | 0.9174 |
| | 40 | 0.8760 | 0.9813 | 0.0187 | 0.0008 | 0.9992 | 0.9257 |
| | 45 | 0.8900 | 0.9829 | 0.0171 | 0.0008 | 0.9992 | 0.9341 |
| | 50 | 0.8944 | 0.9850 | 0.0150 | 0.0009 | 0.9991 | 0.9375 |
| | 55 | 0.8939 | 0.9861 | 0.0139 | 0.0009 | 0.9991 | 0.9377 |
| | 60 | 0.8966 | 0.9869 | 0.0131 | 0.0009 | 0.9991 | 0.9396 |
| | 70 | 0.8737 | 0.9874 | 0.0126 | 0.0008 | 0.9992 | 0.9271 |
| | 50,45,40 | 0.9042 | 0.9834 | 0.0166 | 0.0009 | 0.9991 | 0.9422 |
| GATK | N/A | 0.8543 | 0.9809 | 0.0191 | 0.0008 | 0.9992 | 0.9132 |
| Pilon | N/A | 0.8649 | 0.9910 | 0.0090 | 0.0008 | 0.9992 | 0.9237 |
| Racon | N/A | N/A | N/A | N/A | N/A | N/A | N/A |
| *Genome* | | | | | | | |
| ntEdit | 40 | 0.8352 | 0.9697 | 0.0303 | 0.0009 | 0.9991 | 0.8974 |
| | 45 | 0.8992 | 0.9836 | 0.0164 | 0.0009 | 0.9991 | 0.9396 |
| | 50 | 0.8904 | 0.9824 | 0.0176 | 0.0009 | 0.9991 | 0.9341 |
| | 60 | 0.8998 | 0.9875 | 0.0125 | 0.0009 | 0.9991 | 0.9416 |
| GATK^ | N/A | 0.9207 | 0.9801 | 0.0199 | 0.0010 | 0.9990 | 0.9478 |
| Pilon | N/A | N/A | N/A | N/A | N/A | N/A | N/A |
| Racon | N/A | N/A | N/A | N/A | N/A | N/A | N/A |

*NOTE: Mean over 3 runs is shown for all statistics. Racon stats are unavailable, as it does not output base edit coordinates

^Stats averaged across all chromosomes

#### Supplemental table 7. Suggested ntEdit options for polishing Illumina and SMS* assemblies with Illumina reads

| | ntEdit parameters | | | | |
|---|---|---|---|---|---|
| Genome size | k | i | d | x | y |
| <10 Mbp | 25 | ≤4 | ≤5 | 5 | 9 |
| 10-500 Mbp | 35 | ≤4 | ≤5 | 5 | 9 |
| 100-3000 Mbp | 40-50[1] | ≤3 | ≤3 | 5 | 9 |
| >3 Gbp | 50 | ≤3 | ≤3 | 5 | 9 |

*SMS: Single-molecule sequencing

[1]A lower k (ie. k40) may work best with a more erroneous assembly (see Table S8), and a larger k (ie. k50) with a more accurate one. We recommend running ntEdit iteratively with different k values for best results. We do not recommend setting the indel options (–i and –d) too high on large genomes, especially if they are assembled from low indel Illumina reads

Note: These are suggestions based on tests reported herein. We encourage users the explore a wide range of options, which may work best with their data

#### Supplemental table 8. BUSCO* analysis on NA12878 human assemblies, pre and post polishing with 54X Illumina reads.

| Assembly[1] | Comparator | #Edits (M) | Run time hh:mm[2] | Peak RAM (GB) | Complete BUSCO[3] (%) | Frag-mented BUSCO | Missing BUSCO |
|---|---|---|---|---|---|---|---|
| **PacBio** | Baseline | N/A | N/A | N/A | 5285 (85.4) | 458 | 449 |
| (Pendleton et al., 2015) | ntEdit | | | | | | |
| | k40 i1 d1 | 3.52 | 2:10 | 99.6 | 5650 (91.2) | 261 | 281 |
| N50 = 25.41 Mbp | k40 i2 d2 | 3.61 | 2:10 | 96.8 | 5650 (91.2) | 262 | 280 |
| | k40 i3 d3 | 3.63 | 2:10 | 97.9 | 5651 (91.3) | 263 | 278 |
| | k50 i1 d1 | 3.44 | 2:10 | 87.4 | 5643 (91.1) | 258 | 291 |
| | k50 i2 d2 | 3.53 | 2:10 | 98.2 | 5647 (91.2) | 259 | 286 |
| | k50 i3 d3 | 3.54 | 2:10 | 97.6 | 5647 (91.2) | 258 | 287 |
| | GATK | 2.66 | 42:21 | 38.0 | 5285 (85.4) | 460 | 447 |
| | Racon[4] | N/A | 40:55 | 21.0 | 5670 (91.6) | 243 | 279 |
| **Nanopore** | Baseline | N/A | N/A | N/A | 5647 (91.2) | 274 | 271 |
| (Jain et al., 2018) | ntEdit | | | | | | |
| | k40 i1 d1 | 0.79 | 2:10 | 41.3 | 5662 (91.4) | 262 | 268 |
| N50 = 7.67 Mbp | k40 i2 d2 | 1.13 | 2:10 | 42.5 | 5663 (91.5) | 258 | 271 |
| | k40 i3 d3 | 1.29 | 2:10 | 46.8 | 5669 (91.6) | 256 | 267 |
| | k50 i1 d1 | 0.59 | 2:10 | 40.8 | 5663 (91.5) | 262 | 267 |
| | k50 i2 d2 | 0.84 | 2:10 | 41.9 | 5673 (91.6) | 250 | 269 |
| | k50 i3 d3 | 0.95 | 2:10 | 44.4 | 5670 (91.6) | 250 | 272 |
| | GATK | 0.96 | 41:45 | 38.0 | 5654 (91.3) | 270 | 268 |
| | Racon[4] | N/A | 45:54 | 18.0 | 5681 (91.7) | 234 | 277 |
| **Supernova**[5] | Baseline | N/A | N/A | N/A | 5722 (92.4) | 228 | 242 |
| | ntEdit | | | | | | |
| N50 = 44.06 Mbp | k40 i1 d1 | 0.08 | 2:20 | 133.8 | 5725 (92.5) | 225 | 242 |
| | k40 i2 d2 | 0.08 | 2:18 | 133.9 | 5725 (92.5) | 225 | 242 |
| | k40 i3 d3 | 0.09 | 2:18 | 133.9 | 5725 (92.5) | 225 | 242 |
| | k50 i1 d1 | 0.09 | 2:19 | 134.4 | 5723 (92.4) | 228 | 241 |
| | k50 i2 d2 | 0.09 | 2:18 | 134.4 | 5724 (92.4) | 226 | 242 |
| | k50 i3 d3 | 0.10 | 2:18 | 134.0 | 5724 (92.4) | 226 | 242 |
| | GATK | 0.19 | 45:05 | 39.0 | 5724 (92.4) | 226 | 242 |
| | Racon[4] | N/A | 44:31 | 11.0 | 5722 (92.4) | 224 | 246 |

[1]The NA12878 assemblies are discussed in detail in Watson and Warr (2019). N/A: Not Available.

[2] Run times are indicated for the full tool pipeline (includes read indexing, alignments, Bloom filter creation, where applicable). ntHits ran in 2h08m and 2h15m and required 36.9 and 38.5 GB RAM to process 322.4M Illumina read pairs (2x251bp, ~54X coverage of human genome) at k40 and k50, respectively. The ntEdit runs completed within 2-4 minutes

[3]BUSCO resources: Lineage, euarchontoglires_odb9 (eukaryote), Number of BUSCO searched: 6192

[4]Racon polishing was run in parallel by splitting each assembly into 20 partitions, and polishing each partition separately (see supplemental methods text). Runtimes are summed over all partitions

[5]Assembly from: https://support.10xgenomics.com/de-novo-assembly/datasets/2.1.0/wfu
